# Communicability systematically explains transmission speed in a cortical macro-connectome

**DOI:** 10.1101/117713

**Authors:** Masanori Shimono, Naomichi Hatano

## Abstract

Global dynamics in the brain can be captured using fMRI, MEG, or electrocorticography (ECoG), but models are often restricted by anatomical constraints. Complementary single-/multi-unit recordings have described local fast temporal dynamics. However, because of anatomical constraints, global fast temporal dynamics remain incompletely understood. Therefore, we compared temporal aspects of cross-area propagations of single-unit recordings and ECoG, and investigated their anatomical bases. First, we demonstrated how both evoked and spontaneous ECoGs can accurately predict latencies of single-unit recordings. Next, we estimated the propagation velocity (1.0–1.5 m/s) from brain-wide data and found that it was fairly stable among different conscious levels. We also found that the anatomical topology strongly predicted the latencies. Finally, *Communicability,* a novel graph-theoretic measure, could systematically capture the balance between shorter or longer pathways. These results demonstrate that macro-connectomic perspective is essential for evaluating detailed temporal dynamics in the brain.

**Author Summary:** This study produced four main findings: First, we demonstrated that ECoG signals could predict the timing of evoked electrical spikes of neurons elicited by visual stimuli. Second, we showed that spontaneous ECoG recorded under a blindfold condition (without any stimuli) could also predict the timing of visually evoked neuronal spikes. We also clarified that performance predictions from blindfold data are essentially supported by the constraints of structural paths. Third, we quantified the propagation velocity (conductance velocity) as 1.0–1.5 m/s, and found that the velocity was stable among different conscious levels. Fourth, *Communicability* successfully characterized the relative contributions of shorter and longer paths. This study represents an important contribution to the theoretical understanding of the brain in terms of connectomics, dynamical propagations, and multi-scale architectures.

## Introduction

The brain can be thought of as both a biological and a physical system, in which electrical signals propagate along axonal or dendritic wiring. The propagation pattern eventually emerges as various cognitive functions and internal thoughts. Propagations along underlining connectivity or wiring can be ubiquitously observed in biological networks^14^, the spread of infections^59^, and the organization of the internet^32^. To understand such propagation phenomena, quantitative evaluations that consider the constraints caused by underlying structural networks are critically important. Quantitative evaluations and interpretations have been supported by graph-theory-based approaches^85, 6^. For instance, the comprehensive network (connectomics) approach is essential for studying brain wiring^13^, and graph-theoretic analyses have been used to study a range of relevant topics, such as the Small-World property, which can explain why spatially distant brain regions are able to communicate easily^7^, hubs and rich club organization, which can be used to extract a collection of highly-connected nodes^79^, and community architecture, which can characterize global groups of nodes^73^. The basic concepts of these approaches to network analysis have been previously summarized in textbooks on graph theory^51, 5^.

Preceding graph-theoretic evaluations of detailed topologies, the extent to which structural networks are similar to functional or effective networks^27^, which can be reconstructed from recordings of long-term neuronal activity, is a fundamental question^33^. This issue is also essential for studying microscopic neuronal networks^1, 68, 51, 40^. Recently, connectomics studies have been made possible due to the massive efforts of collaborating teams, and the quality and resolution of data have gradually improved^29,51,55^. The main focus of these studies is often structural networks or spatial patterns of relatively stable neuronal activities^26^. While characterizing relatively stable architecture, studies have gradually emphasized the importance of the dynamics of functional network architectures^39, 25, 54^. However, very few studies have satisfied the following criteria: (1) millisecond temporal resolution, (2) treating the whole-brain as one system, (3) inclusion of structural constraints, and (4) exclusion of computational demands of localized electrical activities.

To address these criteria, we gathered multiple data sets recorded via three modalities: The first modality, ECoG, is a promising technique for capturing the propagation of electrical signals in a large cortical region^44^ or whole cortex^16^. We expected that combining ECoG data with neuronal spike signals would provide a neuronal or microscopic scheme of macroscopic brain signals^52^. As mentioned, we also included structural network data to express the anatomical fibre pathways.

When theoretically testing electrical propagations along a set of pathways, it is possible to simply consider the contributions of the shortest paths, or only the directly connected pathways. However, a recent study demonstrated the importance of considering the non-shortest paths^29^. A graph-theoretic measure, termed Communicability, provides a systematic framework for assessing the balance between shorter and longer paths (walks)^23, 24^. However, Communicability has not yet been used to test the relationships among different neural propagations, which can spread throughout the whole brain within milliseconds.

In the present study, we asked four basic questions regarding the time taken for electrical signals to propagate through the brain: First, focusing on propagations of evoked electrical signals in the primate cortex, we asked how well the global transmissions of electrical signals recorded with ECoG could predict the onset timings of neuronal spikes. Second, to check the robustness of this predictive ability, we evaluated how well time delays in ECoG data could predict time delays in evoked neuronal spikes in the absence of a clear stimulus onset. Third, as a simple but fundamental question, we estimated the propagation velocity of globally propagating electrical signals. Fourth, we examined the possibility of creating fundamental graph-theoretic descriptions of propagation using Communicability.

## Results

### 1. Comparison of evoked activities between ECoG and neuronal spikes

First, we tested how well macroscopically recorded neuronal activities, i.e., ECoG evoked signals, could predict microscopically recorded activities, i.e., spikes in a visual-task condition. In the ECoG data, the transmission delays are given as the time delays of the primary peak in the individual time series of evoked spikes that occurred at the target region within 100 ms of stimulus onset via the primary visual region (Fig. 1-A). Then we evaluated the sharpness of the averaged waveforms using a variable named Peak Index (subsection 3 in Materials and Methods, Figure 1-B) to extract the optimal time window. If the time window is too long, it is possible to erroneously select indirect or separated pairs of brain regions. Equally, if the time window is too short, long connections, which have long propagation times, may be ignored. We expected that responses in brain regions directly connected through anatomical pathways would be sharper than responses in indirectly connected brain regions.

**Figure 1.**
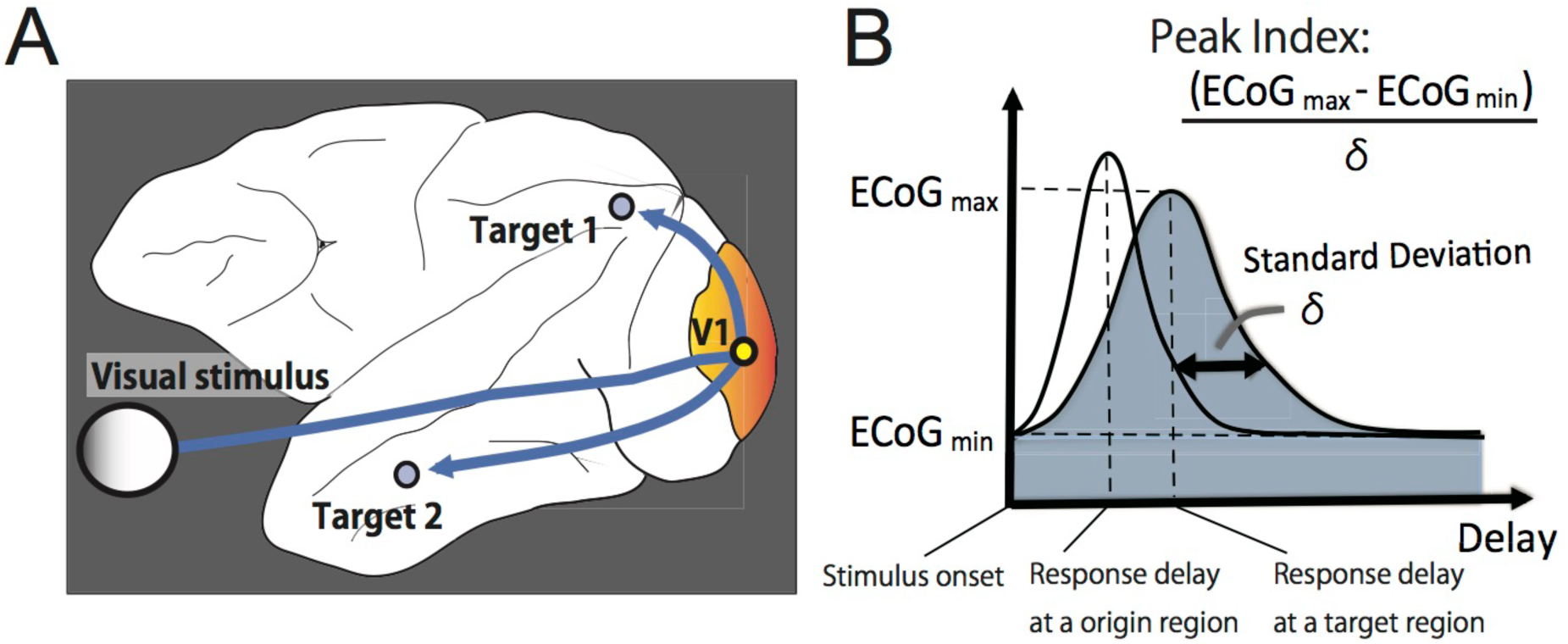
The definition of transmission delays in evoked experiments. A. Schematic illustration showing prominent visually-evoked electrical activity transmitted via the primary visual area (V1), to other “target” brain areas. B. The transmission delays were simply determined as the primary peak of evoked activities after the onset of an individual visual stimulus. To evaluate the sharpness of evoked responses, we defined a variable named *Peak Index* using the equation in panel B. We used the average *Peak Index* for all “target” regions to optimize the size of the time window in which we searched for peak points of evoked activities.

In the spike data, the time delays were given by mode values of firing rates locked to the visual stimulation. The result is shown in Figure 2. The lower panel in Figure 2-A shows the change in the Peak Index depending on the delay after the visual stimulus onset in the ECoG experiments. These coloured maps confirm the presence of a clear flow of electrical activity expanding to whole brain regions, and indicate that the flow is stable against changes in the size of the time window used to search for peak points. However, the consistency of reconstructed electrical flows, quantified by the averaged Peak Index, depended on the size of the time window.

**Figure 2.**
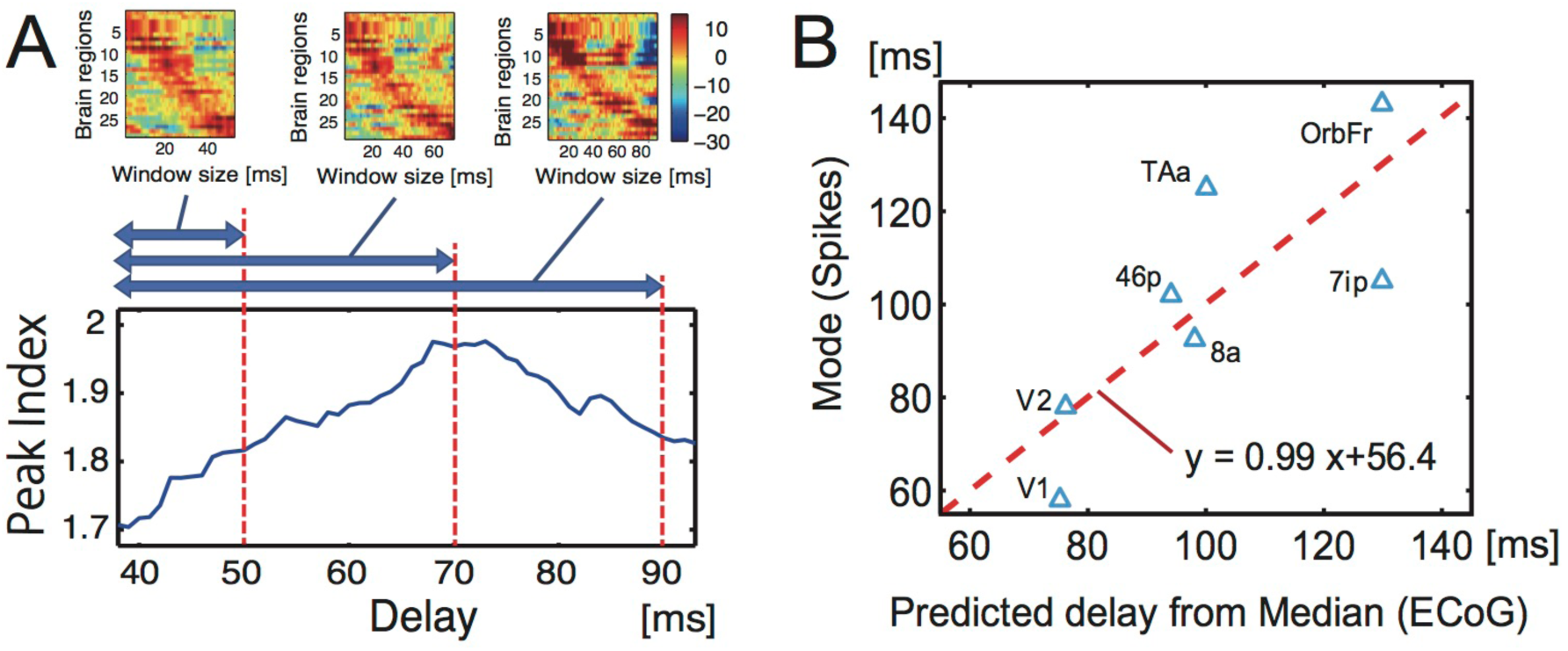
Comparison of delays between neuronal spikes and ECoG within stimulus-driven activities. The bottom section of panel A shows the relationship between the length of the time window used to select peak points of cross correlations (the x-axis) and the averaged *Peak Index* of the waveforms of all electrodes (y-axis). The upper panels in panel A are colour maps of ECoG signals at three different time windows, 0–50, 0–70 and 0–90 ms. The x-axes indicates the size of the time windows and the y-axis is the index of brain regions expressed in terms of ECoG sensors. The values on the y-axis were sorted according to the time delay of the peak point. Panel B shows the results of the main predictions of delays of visually evoked electrical spikes (y-axis) by fitting a linear model to the ECoG signals (x-axis).

The main panel in Figure 2-A shows that the averaged Peak Index was maximized when we searched for peaks in a time window that was 70 ms or shorter. Thus, this time window is optimal for detecting the clearest flow of electrical activity through the macaque cortex. Therefore, we determined the time delays for individual pairs of brain regions in the time window 0–70 ms. Figure 2-B shows the scatter plot of time delays in neuronal spikes (y axis) and predicted delays of spikes from evoked ECoGs (x-axis) according to a linear regression model (y=ax+b). In the regression model, the gradient value was close to 1, indicating that we selected an appropriate time window. Therefore, we searched for activity peaks at 35 (40–75) ms, starting from the primary visual area: A time delay of 40 ms was previously estimated as the time required for visual information to travel from the retina to the primary visual region^38^.

### 2. Comparison between neuronal spikes and ECoG in the task-free condition

So far, we have related visually evoked ECoG dynamics to spike-based latencies evoked by visual stimuli. Next, we sought to determine how well the ECoG signal flow during the Anesthetized and Awake Task-Free conditions, which had no clear visual stimulus onset, could reproduce the time delays recorded as electrical spikes (see subsection 1 in Materials and Methods). The relationship between evoked and spontaneous activity is a fundamental issue in neuroscience^3, 34^. Even in non-stimulated conditions, our brains are always working to process visual and motor information internally^58, 62^.

In the Anesthetized and Awake Task-Free conditions, we estimated the time delays in three steps (Figure 6, subsection 4 in Materials and Methods): First, we adopted the peak delays of cross-correlations between two time series at two brain regions (see subsection 3 in Materials and Methods) as the time delays for signal transmission on a direct pathway connecting the two brain regions. Second, we summed the delays necessary for all individual path components on the pathway. The example in Figure 6 shows pathways connecting region I to region J. Third, we calculated the weighted average of all delays for all pathways based on the three different weight models.

The three “Walk Ensemble Models” (Fig. 3-A), which determine how to add individual time delays along selected chains of edges in three different ways, lead to completely different trends. The chain of edges referred to as a *Walk,* a graph-theoretic concept, is a set of nodes connected successively by links such that connecting back to the same node is allowed. Interestingly, time delays predicted from the Shortest Walk (SW) Model showed a clear positive correlation with spike timings, while the Mean Walk (MW) Model showed a clear negative correlation. The SW Model gives excessively high weights to walk ensembles holding the shortest walks. Inversely, the MW Model gives excessively high weights to ensembles holding longer Walks because the number of samples holding longer Walks is exponentially longer than the number of Walks. Therefore, we designed an intermediate model, the Decay Walk (DW) model, using weights that decay exponentially depending on the increase in Walk *n*. For example, if the transmission probability decreases by *α* (0 < *α* < 1) per one walk step, the multiplied transmission probability for *n* steps of Walk would be expressed as *c_n_ = Prob*(*α,n*) *= α^n^* (0 < *α* < 1). This corresponds with the natural probability expressing how often individual walks will be used in a random walk process. Depending on the index *α*, the DW model gradually changes from behaviour similar to the SW model to that similar to the MW model. If *α* = 1, the DW model corresponds with the MW model, and for the limit *α* → 0, we expect the result to approach that of the SW model.

**Figure 3.**
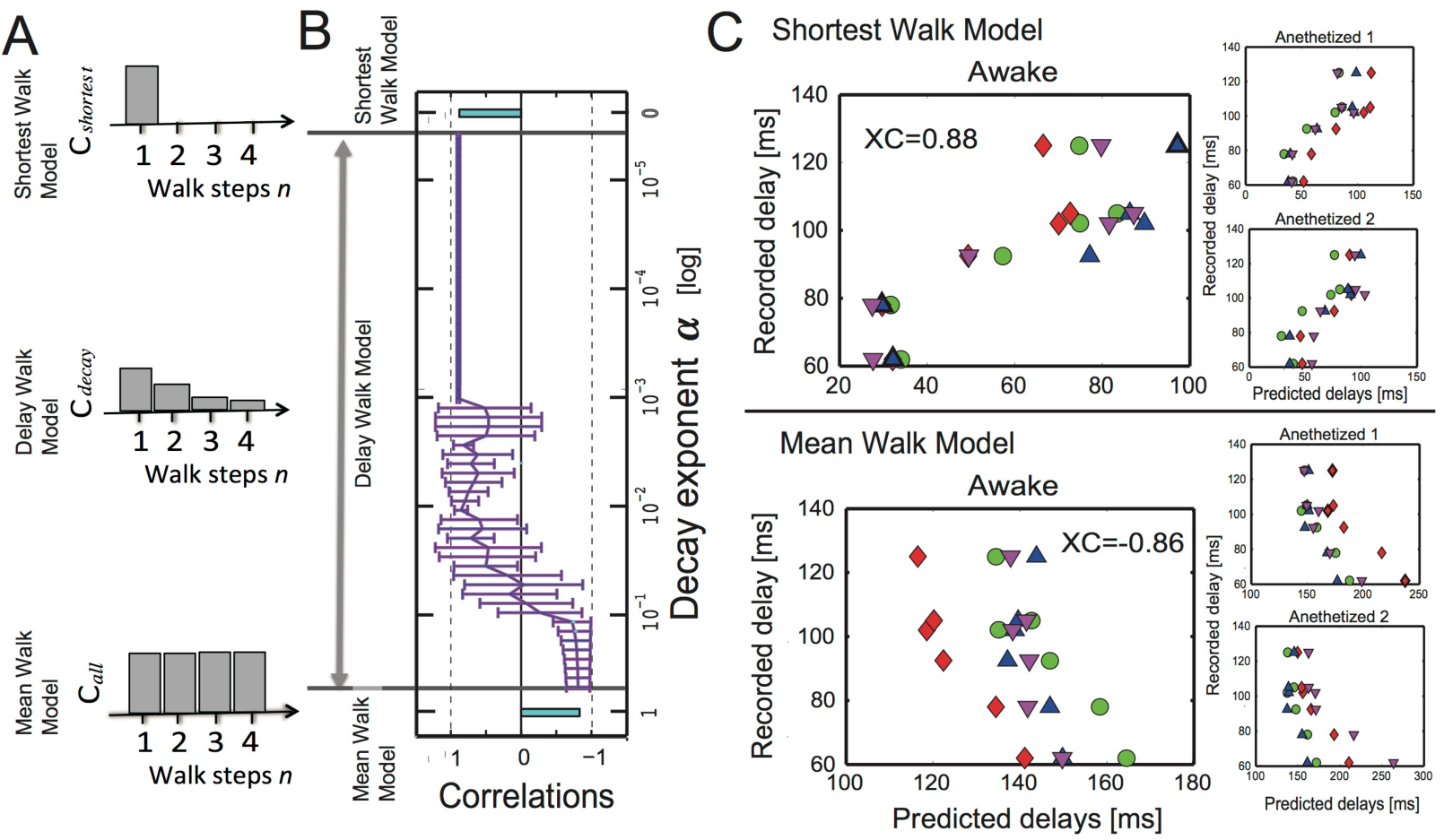
Predictions of time delays of electrical spikes from ECoG data under the no-task condition. Panel A shows histograms of the weights used to prepare the three Walk Ensemble Models. From top to bottom, the histograms show the weights for the Shortest Walk (SW) Model, the Decay Walk (DW) Model, and the Mean Walk (MW) Model. Panel B shows the correlations between delays of neuronal spikes and delays predicted using the three models. The top and bottom bars are correlations for the SW and MW model, respectively. In the MW Model, the mean Walks were calculated from samples included in the one-sigma window (mean value ± one standard variation). The intermediate line on the y axis between the two models corresponds to the correlations for the DW model, determined as a function of exponent *α* in the equation *c_n_* = *α^n^*. In panel C, the upper three scatter plots show the results for data processed with structural constraints based on the SW model, and the lower three panels are scatter plots for the MW model. In both cases, the biggest panels are the results under the awake task-free condition, and the two small panels show the results for the two anesthetized conditions. The four different markers indicate the four different individual monkeys.

Interestingly, we found that at an intermediate *α* in the DW model, the correlation between the spike delays and the delays estimated from spontaneous ECoG under the constraint of structural connectivity, reversed from a strong positive to a strong negative value (Fig. 3-B). The scatter plots between the original spike delays and estimated delays show natural diagonal distributions at strongly correlated regions (Fig. 3-C). At the intermediate phase (10^−2^–10^−1^), the correlation gradually changed between these positive and negative values. This result indicates that the first electrical signals (< 100 ms) to reach their destinations in brain networks use ensembles of shorter paths more often than ensembles of longer paths. It seems that the exponent *n* must be sufficiently smaller than 10^−1^ in the weight *α*^n^. In subsection 4, we will address what may have determined the transition point.

### 3. Time delay for spatial spreading and conduction velocity

So far, we have not observed spatial dimensions. Because brain regions are embodied in space, the spatial coordinates should also reflect the temporal dynamics of propagating electrical signals. Therefore, we examined the relationships between the spatial distances summed along the walk steps from the original to the target brain regions, and the necessary delays for electrical propagations through these walks (Fig. 4).

**Figure 4.**
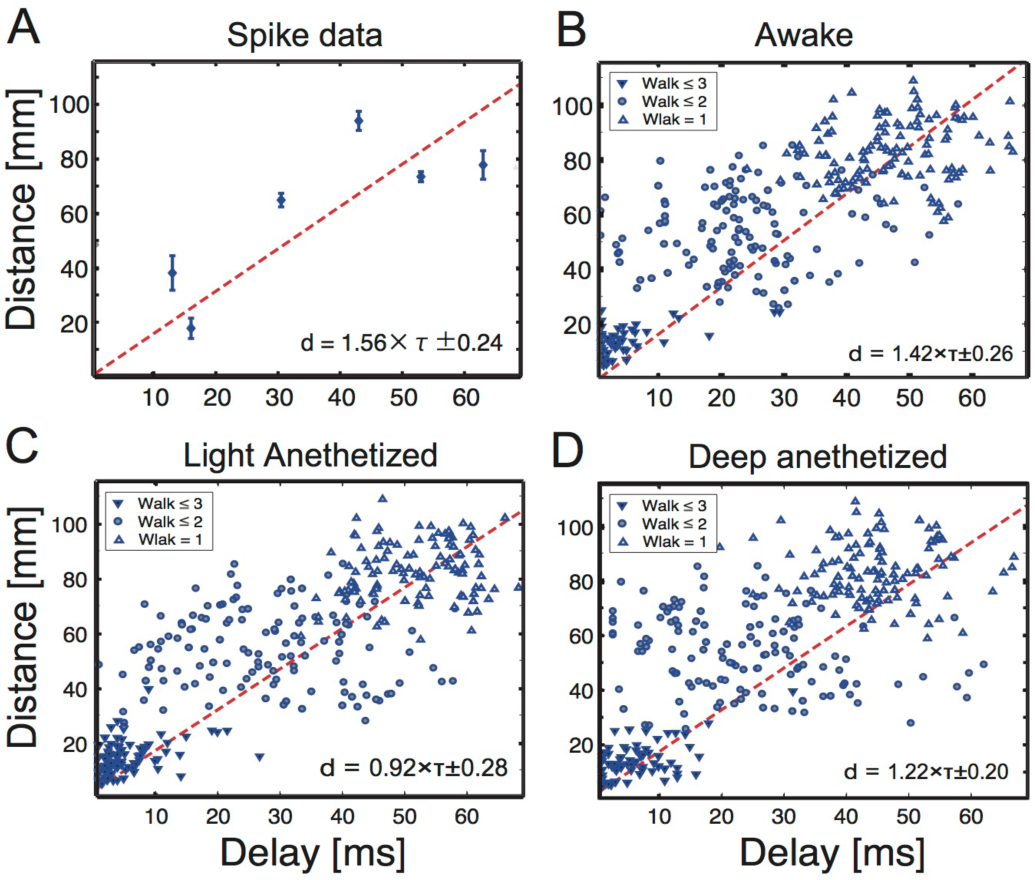
Estimating conduction velocities on the cortical connectome at three conscious levels. Panel A contains a scatter plot showing distances between pairs of brain regions vs necessary delays to transmit neuronal spiking activities between them. Panels B-D show three dense scatter plots of the relationships between distance and necessary delays, estimated from the ECoG data. The three panels B-D reflect data for different cognitive states (awake state, light/deep anesthetized states). In the three panels, the downwards-pointing triangle markers indicate the results for directly connected paths (Steps of Walks *n* = 1), circles denote the results for samples with *n* = 2, and upwards-pointing triangle markers correspond to samples with *n* = 3. The inserted equations in the individual panels are equations for fit lines (d: distance, τ: delay). The two dotted lines in each panel are fit lines for samples with Walks that contain less than 4 steps. When the distance between two brain regions is longer (y-axis), the transmission requires more time (x-axis). In all states, the conduction velocity (the slope of the fit line) was ~1.0–1.5 m/s.

The conduction velocity of electrical brain signals is a fundamental question in neuroscience. Due to limitations in technology or existing data, it has not yet been possible to estimate the velocity from brainwide observations. Here, we estimated the velocities for three conscious or arousal levels as ranging from 1.0–1.5 m/s. Interestingly, the estimated velocity was fairly close to the conduction velocity estimated from the perspective of optimally synchronous brain states in a computational simulation study^20^. We found that the velocities were only slightly different (not significant) among the different conscious levels (p > 0.3, panels B-D in Figure 4).

### 4. Walk ensemble models and Communicability

In section 2, we reported that the time delays in firing spikes could be successfully estimated from ECoG data when we considered the cases in which shorter Walks (with structural constraints) are used more frequently than longer Walks. The balance between shorter and longer Walks seems to be characterized by *α* in the Delay Walk (DW) model. Thus, our final topic in this report involved determining the transition point according to *α.*

The DW model has exactly the same form as a class of novel measure: the Communicability between two nodes of a network. This was introduced in a series of studies of complex networks^23^ [see also equation (15) in Ref. 24]. Importantly, *<u>Communicability can systematically quantify how longer walks contribute to the spread of information in many systems, including the brain</u>.* Let *A* denote the adjacency matrix of the network; each element *A_ij_* is one if a node *i* is linked to a node *j,* and is zero if not. The *Communicability* between nodes *p* and *q* can be defined as the (*p,q*) element of the matrix (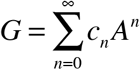). The (*p,q*) element of the summand (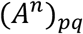) is equal to the number of Walks that connect nodes *p* and *q* in *n* steps. The Communicability *G_pq_* therefore takes account of each *n*-step Walk with the weight *c_n_*. The weight can be *c_n_ = α^n^* with a small parameter *α,* for which *G =* (*I – αA*)^*−*1^, or (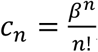), for which *G = e^βA^* To more systematically understand the given results for our neurophysiological data, we also calculated Communicability as a function of the decay factor *α*. Figure 5-B is the correlation between Communicability between a pair of brain regions and the necessary delays required to transmit neuronal spikes between them. For reference, Figure 3-B, which shows the correlation between the necessary delays and the Delay Walk Model, is reproduced as 5-A. The light grey region (_*α*_ 0 07) corresponds with the region where negative correlations were observed, and the negative correlation was reversed at the bottom region (*α* 0 07). We also observed this trend for the correlations with Communicability in all cases when we limited the maximum number of Walk steps to 3–5, although the trend changed when we limited the maximum Walk steps to 2 (Fig. 5-B). This result indicates that at least 3 steps of Walk should be considered to properly characterize the balances between the shorter and longer paths. The original Communicability is also shown as Figure 5-C. At the region where correlations were stable, Communicability also held a stable value.

**Figure 5.**
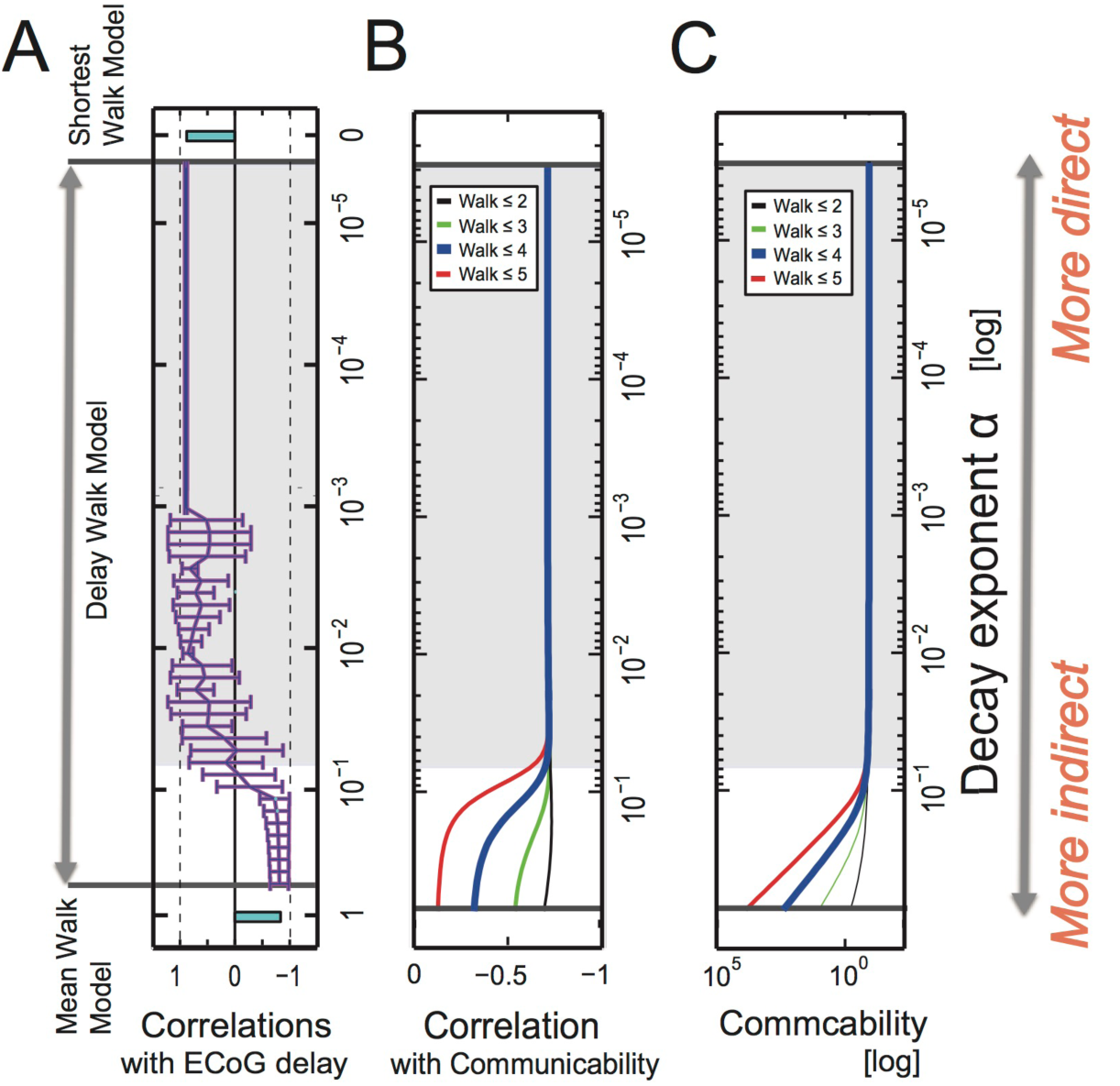
Predictions of time delays of electrical spikes from Communicability. Panel A is the same figure from panel B in Figure 3. The correlation between time delays recorded by electrical spikes and the estimated time delays from the ECoG data is described as a function of the decay factor *α*. Panel B shows the correlations between Communicability and the delays of the electrical spikes. The four lines correspond to cases in which the maximum Walk steps were limited to 2,3,4, and 5 steps. Panel C shows the original Communicability as a function of *α*. The four lines have the same meaning as those shown in panel B.

## Discussion

### 1. Main findings

This study produced four main findings: First, we demonstrated that ECoG signals can be used to predict the timing of evoked electrical neuronal spikes elicited by visual and auditory stimuli. Second, we confirmed that spontaneous ECoG under a blindfold condition (without stimuli triggers), can predict the timing of visually evoked neuronal spikes. The prediction performance from the blindfold data was efficiently supported by structural constraints. Third, we estimated the propagation velocity (conductance velocity) as 1.0–1.5 m/s using connectomic data, and found that the velocity does not depend on conscious level. Fourth, we demonstrated that Communicability can be used to systematically characterize the contributions of the shortest paths and non-shortest paths in the general pattern of transmission delay.

### 2. Multi-scale neuronal recording technologies

We were able to predict the time delays of visually evoked spikes from ECoG data recorded in the blindfold condition. Previous studies have successfully predicted spatial patterns of functional spontaneous activities observed from fMRI using spatial patterns of structural networks from diffusion tensor/spectral Imaging^33,21,30^. The temporal resolution of fMRI is longer than one second. To contrast this, we aimed to show how high structural constraints could influence the determination of temporal dynamics of neuronal spikes using a higher temporal resolution signal, i.e., ECoG, which is less than a millisecond. Several studies have reported high prediction performance of spike timings from Local Field potentials (LFPs)^60,42,61^. Because the spatial scale recorded in ECoG, 1 cm, is over 100 times the spatial scale recoded in LFP, 1 mm, and many complex spatial patterns can be generated in the spatial map, the success of prediction using ECoG signals is a non-trivial result. Additionally, past predictions using LFP tested limited brain regions, while the data set used in our study included whole cortical networks. Therefore, the results of the present study provide important insight into the integration of different spatial scales^66^ and the prominent nonuniformity of the cortical regions observed.

### 3. Subcortical contributions

When simultaneously observing many brain regions, it is important to consider the important roles that subcortical regions play in mediating electrical interactions between cortical regions^64;49^. Indeed, how cortico-subcortico-cortical connections and subcortical pathways influence global dynamics within the cortex is an interesting question for future research^70^. Collaborative studies involving simultaneous recordings from many subcortical and cortical regions will improve our current understanding of neuronal signal transmission^47^.

### 4. Transmission delay and Communicability

The constraints of structural networks on transmission delays may be systematically captured by Communicability. Communicability characterizes the relative contribution of shorter and longer paths. In general, as the path length between two nodes increases, Communicability decreases. Because ECoG delay qualitatively reflects the path length, Communicability showed clear negative correlations at the regions where *α* is small. This corresponds to the Delay Walk model, which showed a clear positive correlation with time delays. In the regions where *α* is bigger, both clear correlations were modulated simultaneously. We also expect slow components, such as P30 0^67, 71^, to reflect activity where the non-shortest paths are more frequently selected, such that a larger *α* will have a stronger contribution.

Communicability has been previously applied to weighted brain data collected via diffusion tensor imaging^17^. Furthermore, Communicability has been found to be a sensitive measure for quantifying changes in brain regions remote from Stroke foci in both an experimental study^18^ and a computational simulation^4^. The removal of nodes with high Communicability, as well as the removal of rich-club nodes, can severely impact global communication in the brain^22^. Interestingly, we found that Communicability could capture the general trend of transmission delays, i.e., the “when” in the macro-connectome.

### 5. Estimation of transmission velocity

We also estimated propagation velocity. With respect to past studies, our main novel contribution is that we evaluated propagation velocity on a global brain scale. Indeed, past studies estimated propagation velocity of neuronal spikes within limited brain regions^75,76,81,83^. In a previous computational modelling study, propagation velocity was estimated in terms of the optimality of synchronous activations between brain regions^22^. In our study, the propagation velocity was fairly stable, even at different Conscious levels. Note that the variance or higher statistical moments of the velocity could potentially describe the differences between the Conscious levels (Fig.4). Meanwhile, we expect that Transfer Entropy will improve the current functional connectivity results better than Cross Correlation, as it will also clarify the relationships between functional connectivity and structural connectivity^30, 68^.

The estimated velocity contained clear variability, and the general form of the histogram of the propagation velocity followed a gamma distribution^53^. A physiological interpretation of the histogram form is a potential topic for future studies. Anatomical connectivity also contains variability in terms of connection strength, and recent studies have reported that there are more weak connections than previously presumed^46^. Physiologically, axonal conduction delays can vary widely depending on myelination or demyelination^81^, axon diameters^83^, and the density of sodium channels^83^, and also depend on the forms of dendritic branches and cell types. Future work evaluating the variety of global propagations^56^ and considering the detailed synaptic topologies of neurons^82,36,78^ will contribute greatly to this field.

### 6. Final remarks

This study focused on the time-delay in cortical information flow. Using Communicability and the index *α,* we quantitatively evaluated how a balance between shorter and longer paths influences the information flow in the unified theoretical framework. How the human brain evolved such an efficient network organization with the selective use of shorter paths remains an interesting question. Shorter paths reduce wiring cost, while some long paths are unavoidably necessary for the integration of information. Therefore, it is important to consider both the optimality and efficiency of the brain structure. We believe that our results represent an important step in generating increasingly realistic predictions of brain dynamics.

## Materials and Methods

### 1. Data acquisition

Using a neuroinformatic approach, we combined three data sets acquired using different modalities by independent research groups: (1) spike-based visual responses in single-unit recordings, (2) brain-wide field dynamics recorded with ECoG, and (3) anatomical connectivity network data among cortical regions from tracer injection studies. All data were collected from the macaque cortex, and processed using the following methods:

**First**, we prepared a summary of responding peak latencies of neuronal spikes from past neurophysiological studies. We assessed neuronal spike timings associated with visual information processes for not only occipital visual areas, including V1 (primary visual) and V2 areas^65^, but also temporal areas such as TPO and TAa^8^, parietal areas including area 7ip^15^, frontal areas such as areas 8a, 46^15,35^, and the orbitofrontal region^77^. Several previous articles have reviewed trends in the time delays of visual evoked activities^38, 12^. The peak latencies of neuronal spikes were represented by the mode values of firing pattern histograms. Note that, although many other studies have recorded evoked firing activities, we limited the data sets included in this study to those that recorded from the cortical gyri. This is because the ECoG data, which will be compared later, was recorded only from gyri. Additionally, if we could not extract mode values (peak points) from figures given in past reports, we excluded that data from our analysis.

**Second,** to optimally model the transmission pathways of electrical signals between brain regions, we considered the constraints of underling structural networks. We prepared the structural network of the monkey brain based on the data given in Lewis and Van Essen (2000)^40^. In their model, the network covers entire cortical regions, and includes the strengths of connections, discretized into seven levels. This atlas is shared publicly in the CoCoMac database^74, 37, 9^. This database has been used in many past studies, and has contributed to many investigations, including a comparison between monkey and human brains^31^, assessment of the relationship between structure and function^11^, and relationship between network structure and cognition^45^. Currently, this database is continuously maintained as the Scalable Brain Atlas^10^.

**Third**, we obtained macroscopic functional data, specifically ECoG data from four macaque monkeys, from the Neurochyco database^16, 48, 80^. The data set includes data recorded continuously from monkeys that were blindfolded and not engaged in any specific tasks, i.e., the “Awake Task-Free condition”. ECoG recordings from anesthetized monkeys are referred to as those collected in the “Anesthetized condition”. The data set also included a visual stimulation experiment. In the visual experiment, a grating stimulus was presented around a fixation cross with one of eight randomly selected grating orientations. The stimuli were shown for 2 s. Refer to the web page (http://wiki.neurotycho.org/Anesthesia_Task_Details) for more detailed information about the ECoG experimental procedure.

### 2. Integration of data

To transform the original structural network data into a network with the spatial resolution of the ECoG sensors, we labelled groups of ECoG sensors according to brain region by comparing photos of ECoG sensors provided in the Neurochyco database with a spatial segmentation scheme of the monkey cortex^41^. Table 1 shows the list of sets of 128 ECoG sensors indexes, the region names for the neuronal spikes, and the indexes in the structural segmentation for individual monkeys. Because the locations of the ECoG sensors were different among monkeys, the corresponding structural brain regions also varied (Right four columns in Table 1). Here, the cortical regions at the sulci or on the longitudinal fissure were eliminated because the ECoG sensors were not indwelled at those regions, which are indicated in the Table 1 as index “0”. The names of the structural brain regions corresponding with the indexes are also separately summarized in Table 2. A previous study used a similar comparison method^67^. Because several regions were eliminated in this process, we regarded pairs of nodes, connected through one intermediate node, as connected. This preprocessing improved the prediction performance of spike timings of neurons^30^.

**Table 1:**
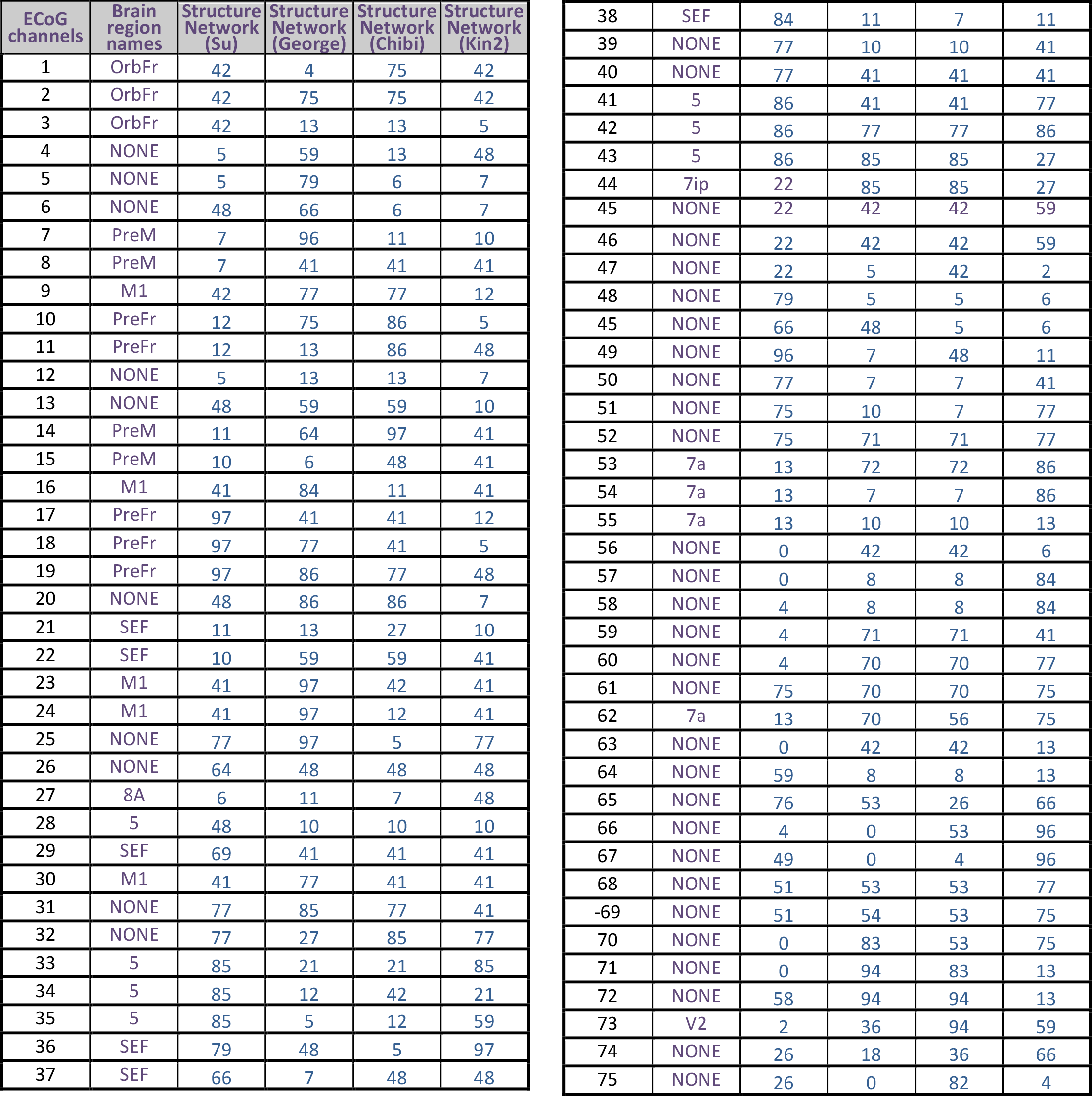
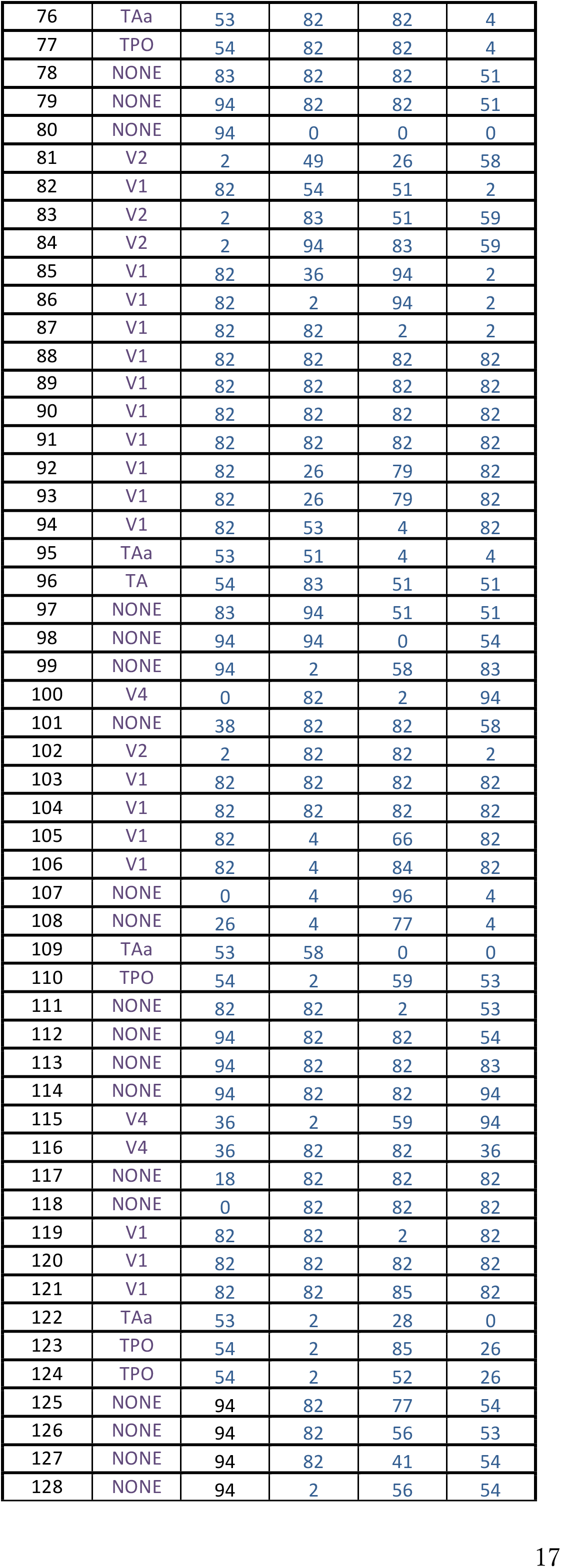
Summary of brain region labels in four monkeys. From left to right, the first column shows ECoG channels from the Neurochyco database. The second column lists the abbreviated names of brain regions that could be used to assess neuronal spikes. Refer to the corresponding references in subsection 1 (‘Data Acquisition’) in the Materials and Methods section. The third–sixth columns contain indexes of structural brain regions located under the ECoG sensors. The names corresponding to individual indexes are summarized in Table 2. Because the locations of the ECoG sensors were different among individual monkeys, the structural regions vary. (OrbFr: Orbital prefrontal cortex, PreM: Pre-Motor Cortex, M1: Primary Motor Cortex, SEF: Supplementary eye field, 5: Area 5, 8A: Prefrontal area 8A, 7ip: Parietal area 7ip, TA: Temporal anterior region, TAa: Area TAa, TPO: Temporal parietal occipital, V4: Visual area 4, V2: Visual area 2, V1: Primary visual area)

**Table 2:**
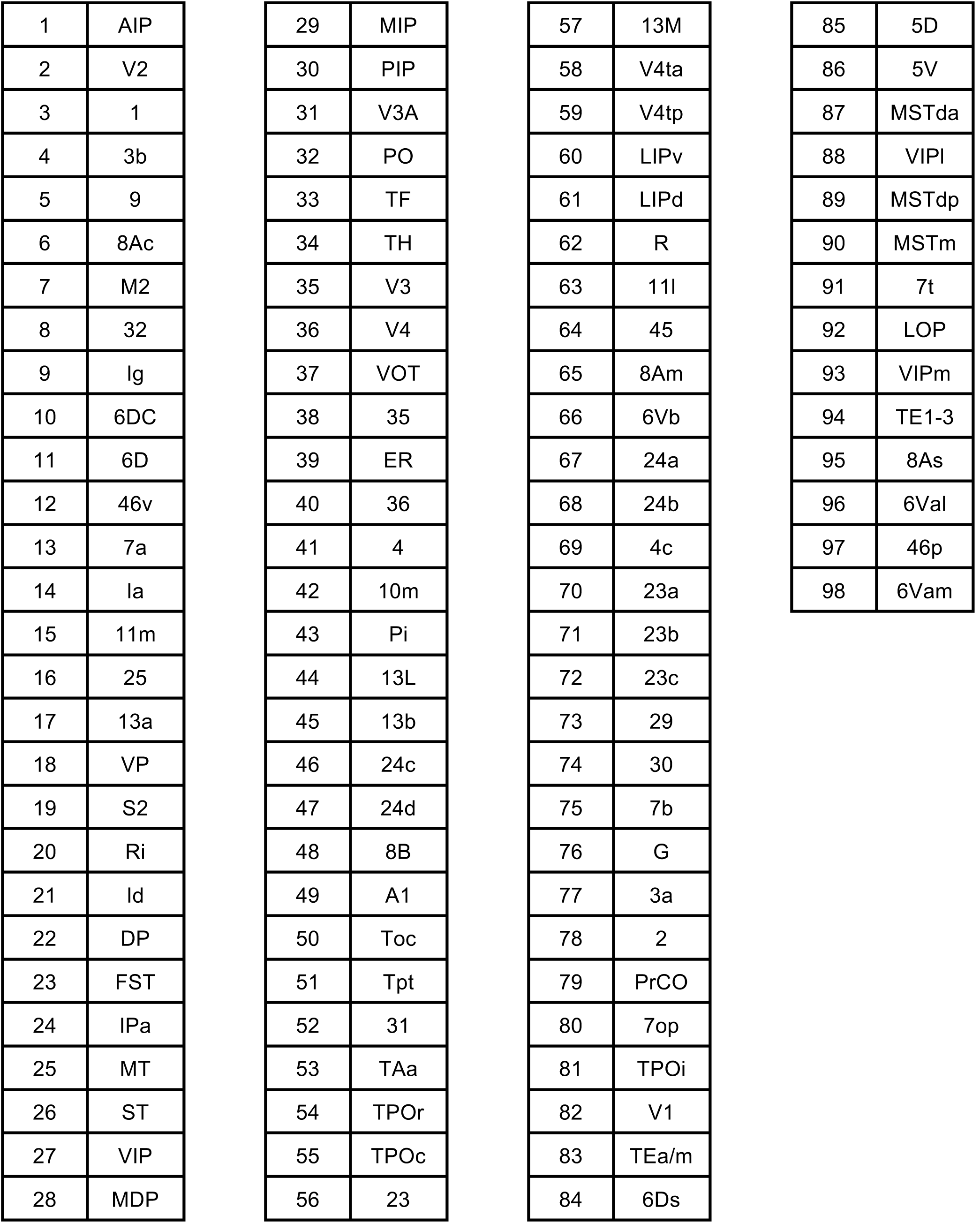
Pairwise list of indexes and corresponding brain regions from Lewis and Van Essen (2000). This table shows the brain regions (right column) corresponding to the indexes shown in Table 1 (left column). Please refer to p.113 in Lewis and Van Essen (2000) for the abbreviation key.

### 3. Estimation of time delays of neuronal spikes from visual stimulus-evoked ECoG activities

In the visual stimulation experiment, we estimated the transmission delay from visual stimulation based on the time of the primary big sharp peak of evoked potential.

As preprocessing, we averaged the 210 trial data points after subtracting the 50 Hz component using a notch (band cut) filter with a 5 Hz standard deviation. Then, we selected the largest peak point between 0- Tms (T < 100ms) after the stimulus onset (Fig. 5-B), and used the time delay of that peak point as the delay of the ECoG evoked data. We explain how the time delays for the anesthetized conditions were extracted in subsection 3 in this method section. Then, we evaluated the sharpness of the averaged waveforms using a variable named *Peak Index* (Fig. 1-B) to extract the most optimal time window. The Peak Index was mathematically defined by the following equation:

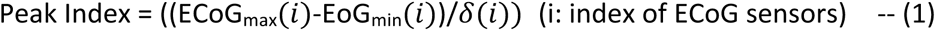

Here, ECoG_max_(*i*) and ECoG_min_(*i*) are the maximum and minimum values, respectively, of the ECoG signal recorded by sensor i, and *δ*(*i*) is the standard deviation when we fit the data to a Gaussian function around the primary peak point. Therefore, this index evaluates the average of the amplitudes at the ECoG peaks for all sensors with sigma *δ*(*i*) as the unit. This value was averaged for all sensors involved in individual brain regions to get an averaged Peak Index representing interactions between brain regions.

### Estimation of time delays from non-time locked ECoG activities alongstructural paths

Here, we explain how we estimated time delays in the absence of a clear stimulus onset, such as in the “Awake Task-Free condition” or the “Anesthetized condition”. This process had three steps: First, instead of evoked activity, we calculated Cross Correlations between all pairs of brain regions, and defined the time delays from the peak forms in the Cross-Correlograms. Second, we identified all possible pairs of ECoG sensors located in the *Origin and Target regions*. Third, we obtained the weighted averages of the delays for all possible pairs of ECoG sensors located in anatomically connected brain regions based on structural networks.

In the first step, after subtracting the 50 Hz components using the same notch filter as that used for evoked activities, we subtracted cross correlations of smoothed components by the following equation:

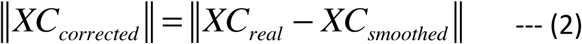

*XC_smoothed_* was calculated by smoothing individual waveforms using a uniform 50 ms time window. From the amplitude of the corrected cross correlations, we detected the primary peak for each pair of Origin and Target regions, within 0–30 ms. We used the time delay at the peak point to characterize the transmission delays of the electrical signals from the “origin” to the “target” region under the Awake task-free and the Anesthetized conditions.

In the second step, we integrated the time delays for the individual pairs of brain regions based on the constraints of the structural network, to predict the time-delays of neuronal firing. More specifically, we searched all pathways connecting all combinations of ECoG sensors between the *Origin* and *Target* regions. For example, as shown Figure 6-A, we selected all ECoG sensors included in regions I and J. We call these i_1_, i_2_, i_3_,…, i_n_ and j_1_, j_2_, j_3_,…, j_m_ respectively. If a pathway from region i_2_ to region j_3_ passes through regions k and I, then we summed the time delays for the three paths: from i_1_ to k, from k to I, and from I to j_3_.

**Figure 6.**
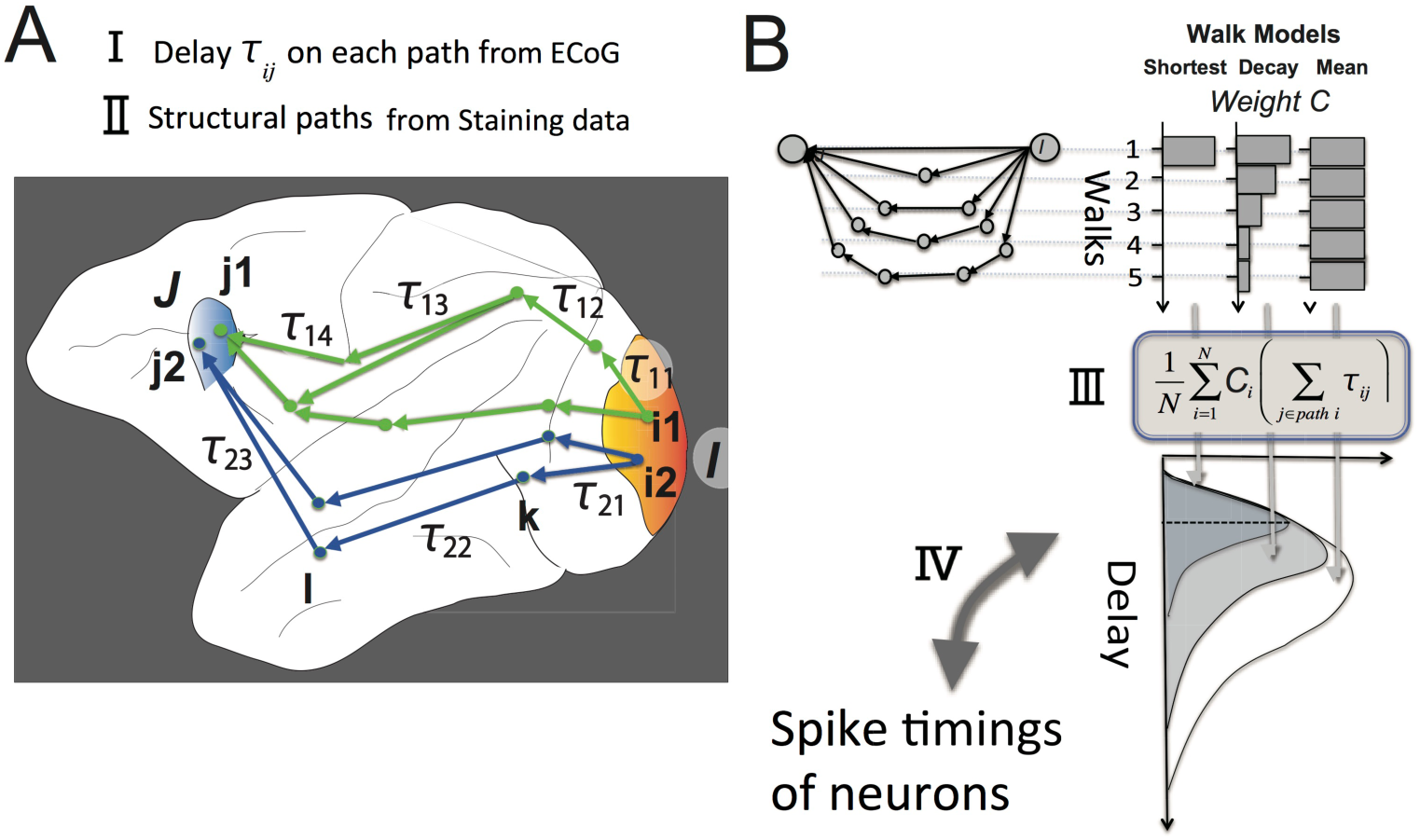
Predictions of time delays of electrical spikes from ECoG data in the Task-Free condition. A. Our scheme involves calculating the time delays from a starting region I to a goal region J using spontaneous activities and structural pathways. In the present study, we used the Cortical Parcellation scheme according to Lewis and Van Essen (2000). Regions I and J are shown as one of the pairs of 98 cortical regions included in the parcellation scheme. There are several ECoG sensors on both regions I and J. Here, we simply consider only two sensors, i1 and i2 (j1 and j2), to exist in each region I (J). In this example, the pathways from region I to region J involve all four combinations of paths from sensors on region *I*, i1 or i2, to sensors on region J, j1 or j2. Each combination of the starting and goal points may involve many paths (Steps of Walk n ≤ 4). Each delay *τ_ij_*_*i,j*_ at a step j on a path i was given as the peak delay of the equation (1) within T ms (T < 100). In panel A, for example, if the activity is transmitted using the most dorsal pathway, the total delay is (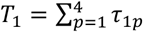) and if the activity is transmitted on the most ventral pathway, the total delay is (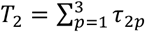) B. We made combinations of possible structural pathways and time delays for each path component. We then prepared three models to calculate the weighted averages of time delays (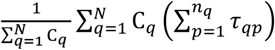) given from different pathways connecting region I to region J. Here, *τ_qp_* is the delay necessary to transmit electrical activities on a pathway connecting node q to node p, and C_*q*_ indicates the relative weights on ensembles with individual Walks. This weight is expressed using three models: The three bar graphs in the top-right show the relative weights for individual Walks for the three models. From left to right, the bar graphs correspond to the Shortest Walk (SW) model, Decay Walk (DW) model, and Mean Walk (MW) model. The individual model provided a different histogram of the sample number of net time delays for the individual related pathway, as shown at the bottom figure in panel B, and the averaged time delays were compared with the time delays given from neuronal spikes data.

Finally, for all detected pathways (Walk < 4)^73^, we calculated the weighted average of the specific time delay for each pair. We defined the weighted average of time delays by the following equation:

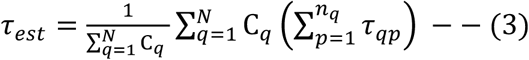

Here, q is the index of the pathway connecting region I to region J, p is the number of Walk steps for a pathway q, and *n_q_* is the maximum number of Walk steps. Therefore, the difference between the weights included in three Walk Ensemble models is reflected in C_*q*_ (Fig. 6-B). The Shortest Walk (SW) model considers only the time delays for the shortest Walks: (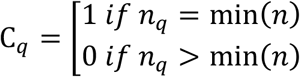) (n: number of Walks). The Mean Walk (MW) model considers all Walks equally, so that C_*q*_ = 1. The Decay Walk (DW) model assigns higher weights to shorter vs longer pathways using the exponentially decaying function C_*q*_ = *α^n^*, where the decay of the exponent of *α* reflects the expectation that longer paths may be used to transmit activities less frequently than shorter paths. When *α* = 1 corresponds with the MW model, and *α* decreases toward 0, the result gradually approaches that obtained using the SW model.

Using these weighted averages (eq. 3), we obtained the representative delays given by the ECoG data under structural constraints. We compared the delays obtained from the ECoG data with the neuronal spike data reported in previous neurophysiological studies (Fig. 6). Notice that the ECoG data had four variations related to arousal level in the Awake condition, and the light/deep Anesthetized conditions. All analyses were performed using Matlab software (The Mathworks Inc.).

## Acknowledgements

MS is grateful to S. Tajima for helpful discussions regarding the properties of ECoG data. MA is also grateful for support received from colleagues and the URA office at Osaka University during this project. This study was supported by a Grant-in-Aid for Research Activity Start-up to MS from MEXT (The Ministry of Education, Culture, Sports, Science and Technology).

## Author contributions

M.S. designed the study, M.S. and H.N. analyzed the data, M.S. wrote the manuscript.

